# Identification of unique α4 chain structure and conserved anti-angiogenic activity of α3NC1 type IV collagen in zebrafish

**DOI:** 10.1101/2022.08.14.503918

**Authors:** Valerie S. LeBleu, Jianli Dai, Susan Tsutakawa, Brian A. MacDonald, Joseph L. Alge, Malin Sund, Liang Xie, Hikaru Sugimoto, John Tainer, Leonard I. Zon, Raghu Kalluri

**Author notes:** co-first authors (Valerie S. LeBleu, Jianli Dai). To whom correspondence should be addressed: Raghu Kalluri: Department of Cancer Biology, Metastasis Research Center, University of Texas MD Anderson Cancer Center, Houston, TX 77054; Valerie LeBleu: Feinberg School of Medicine & Kellogg School of Management, Northwestern University, Chicago, IL 60611.

## Abstract

Type IV collagen is an abundant component of basement membranes in all multicellular species and is essential for the extracellular scaffold supporting tissue architecture and function. Lower organisms typically have two type IV collagen genes, encoding α1 and α2 chains, in contrast with the six genes in humans, encoding α1 to α6 chains. The α chains assemble into trimeric protomers, the building blocks of the type IV collagen network. The detailed evolutionary conservation of type IV collagen network remains to be studied. We report on the molecular evolution of type IV collagen genes. The zebrafish α4 non-collagenous (NC1) domain, in contrast with its human ortholog, contains an additional cysteine residue and lacks the M93 and K211 residues involved in sulfilimine bond formation between adjacent protomers. This may alter α4 chain interactions with other α chains, as supported by temporal and anatomic expression patterns of collagen IV chains during zebrafish development. Despite the divergence between zebrafish and human α3 NC1 domain (endogenous angiogenesis inhibitor, Tumstatin), the zebrafish α3 NC1 domain exhibits conserved anti-angiogenic activity in human endothelial cells. Our work supports type IV collagen is largely conserved between zebrafish and humans, with a possible difference involving the α4 chain.

## Introduction

Type IV collagen (collagen IV) network assembly is essential for basement membrane organization and crucial for multicellular life [1–6]. The formation of the type IV collagen network is a complex process that involves both intracellular and extracellular steps. In the endoplasmic reticulum, three collagen IV α chains assemble into heterotrimeric helical molecules, known as protomers, which serve as the building blocks of the type IV collagen network [7]. After being secreted from cells, type IV collagen protomers undergo oligomerization and self-assembly into insoluble sheet-like supramolecular networks [8–11]. Lower organisms, such as *C. elegans* and *Drosophila*, have two distinct chains, namely α1 and α2, whereas mammals express six distinct chains named α1 through α6, encoded by six distinct genes (*Col4a1 – Col4a6*) [12]. The chain composition of type IV collagen network is in part dictated by unique type IV collagen protomer nucleation and formation, which are specified by α chain non-collagenous (NC1) domain interaction [13–15].

Despite many possible combinations, predominantly only three distinct protomers α1α2α1, α3α4α5, and α5α6α5 have been identified in basement membranes [16, 17]. The collagen IV α1α2α1 network, considered to be more primordial, is ubiquitously found in basement membranes of all tissues. The α1α2α1 network is thought to be critical to the evolution of multicellular life [18–20]. Indeed, genetic deletion of collagen IV α1 and/or α2 genes in mice results in a loss of basement membrane integrity and embryonic lethality [21–23], and loss of the collagen IV α1 alone in *C. elegans* and *Drosophila* leads to developmental arrest and embryonic lethality [1, 24–26]. It is also reported that collagen IV α1 and α2 mutations correlate with diseases in humans [27]. In contrast, collagen IV α3-α6 chains, only found in higher-order organisms, have restricted tissue distribution during development, and present in more specialized basement membranes [17]. For instance, the α3α4α5 (IV) network plays a prominent role in the mammalian kidney, where it is a critical component of the filtration barrier in the glomerular basement membrane (GBM). Mutations in α3-, α4-, and α5-chain encoding genes leads to the absence of α3α4α5 (IV) protomer formation in GBM, which is associated with several nephropathies, including Alport syndrome, thin basement membrane nephropathy, and focal segmental glomerulosclerosis [2, 28–31]. The α3α4α5 protomer is also noted in the basement membranes of many other tissues, including brain, alveoli, testes, cochlea, the synapses of neuromuscular junctions, and lung epithelia in mammals [32–38]. Genetic disruption of the α3α4α5 network composition in these basement membranes is associated with multiple organ impairment in Alport syndrome [39]. In addition, autoantibodies against the NC1 domain of the collagen IV α3 chain results in Goodpasture syndrome, which is characterized by pulmonary hemorrhage and glomerulonephritis [40, 41].

Beyond its likely role in structural stability of Collagen IV network in basement membranes[13], the cleavage of the NC1 domain releases the bioactive α3 NC1 fragment, Tumstatin, which exerts anti-angiogenic and pro-apoptotic functions that significantly influences tumor growth [42–44]. Interestingly, zebrafish, like mammals, express six collagen IV α chains [45]. The development of the collagen IV α3α4α5 and α5α6α5 networks are thought to result from more recent evolutionary events, and these networks may not be as highly conserved as the primordial α1α2α1 (IV) network. Indeed, the α3-chain cryptic epitope/s targeted by the autoantibodies in Goodpasture syndrome do not appear to be conserved in zebrafish [46]. Little is otherwise known about the evolutionary divergence of collagen IV basement membrane composition from the zebrafish to mammals [47, 48]. Here we report on the degree of homology of NC1 domain between the six zebrafish collagen IV α chains and their human homologues. We describe temporospatial expression pattern of type collagen IV chains in zebrafish development at the mRNA level. We also demonstrate that the zebrafish α3 chain NC1 domain (zTumstatin) is present with conserved anti-angiogenic activity when compared to human Tumstatin.

## Results

### The zebrafish type IV collagen alpha 4 chain is distinct from its human homolog

Type IV collagen NC1 domains in mammals are critical for protomer assembly and present with anti-angiogenic activities [49–52], and the head-to-head organization of the Col4α1-Col4α2 pairs, Col4α3-Col4α pairs and Col4α5-Col4α6 pairs was found in human, mouse and zebrafish species (**Fig. 1a**). Each gene encodes for a protein with a characteristic N-terminal 7S domain, a collagenous domain, and a C-terminal non-collagenous domain (NC1) (**Fig. 1b, Fig. 2**). A high degree of amino acid sequence homology between the NC1 domains of zebrafish α1, α2, α5, and α6 chains and the mouse and human orthologs was noted, with more pronounced dissimilarity for α3 and α4 chains (**Fig. 1c, Fig. 2**). Comparative analysis of the zebrafish NC1 subdomains across the six Col4 chains revealed that specific features, critical for the maintenance of the tertiary structure of the NC1 domain and protomer assembly, are highly conserved between zebrafish and human (**Fig. 3a**). All six zebrafish Col4 chains contain the HSQ tripeptides that signal the beginning of the NC1 subdomain I and II (**Fig. 3a**, cyan). Each zebrafish α chain also contains twelve conserved cysteine residues (**Fig. 3a**, green) that maintain the domain structure essential for protomer assembly. It is reported that the collagen IV network is stabilized by protomer dimerization at adjoining NC1 domains, preceded by a chloride-dependent NC1 domain conformational change through salt bridges formed between the R76, D78, E175-R179, and N187 residues [9]. These residues are mostly conserved in all six zebrafish α chains, except for R179 in the α2 chain, for which a R179K substitution in is noted (**Fig. 3a**, red arrow). In addition, the zebrafish α3 and α4 NC1 domains lack the N187 residue (**Fig. 3a**, red). It is speculated that once the NC1 domain alignment occurs, protomers are linked by a sulfilimine bond between the M93 and K211 residues in a reaction catalyzed by the highly conserved peroxidasin enzyme [10, 11, 53]. Five of the six zebrafish α chains (except for the α4 chain) contain the M93 and K211 residues associated with the formation of the sulfilimine bond and protomer dimerization (**Fig. 3a**, yellow) [10].

**Figure 1:**
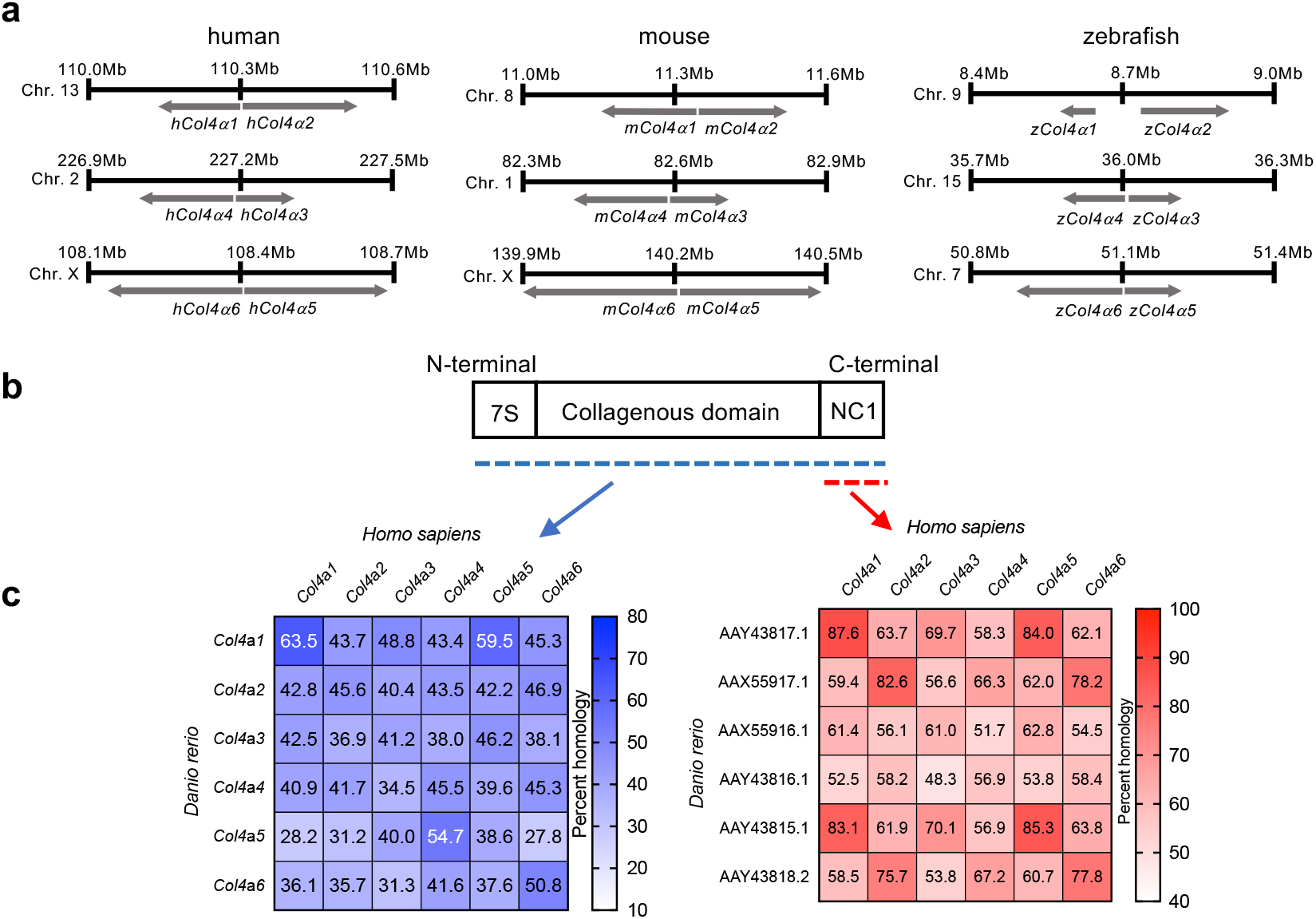
Chromosome mapping for α chains between human, mouse, and zebrafish and amino acid sequence homology. **a**. Chromosome mapping of human, mouse, and zebrafish collagen IV α chains. **b**. Schematic representation of the domains of Collagen IV chains, and **c**. percent homology of amino acids in full length collagen IV (blue) or NC1 domains (red).

**Figure 2:**
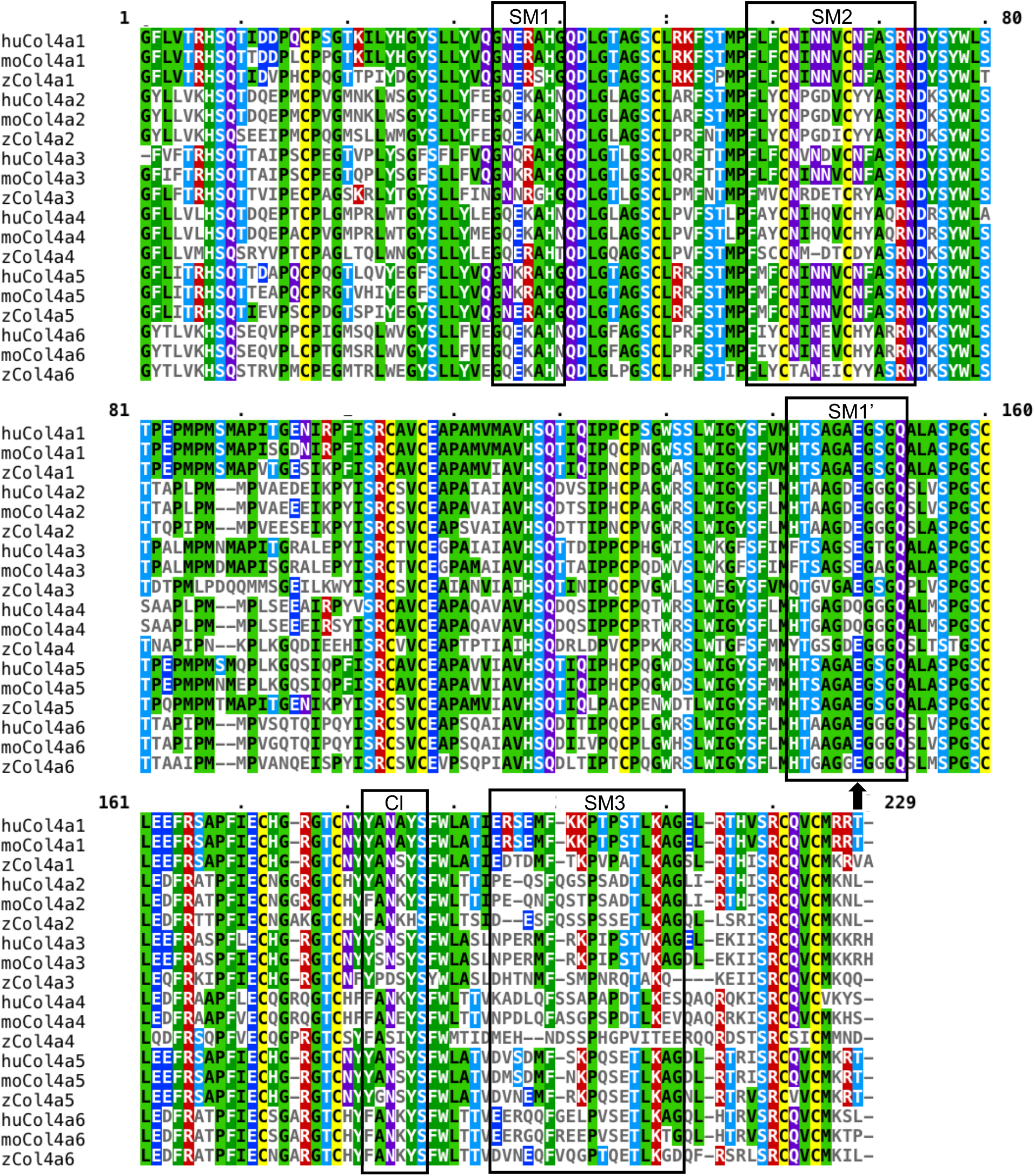
Sequence comparison of the human, mouse, and zebrafish NC1 domain. Comparison of amino acid sequence shows conservation among different species. Subdomains of interests are boxed, structural motifs SM1 (hairpin 3–4), SM2 (hairpin 6–7), SM1’ (hairpin 3’–4’), SM3 (9 and preceding loop), CI (chloride ion). Arrow points to amino acid residue similar in zCol4a4 and hCol4a1 and hCol4a2 NC1 domain. hu, human; mo, mouse; z, zebrafish.

**Figure 3:**
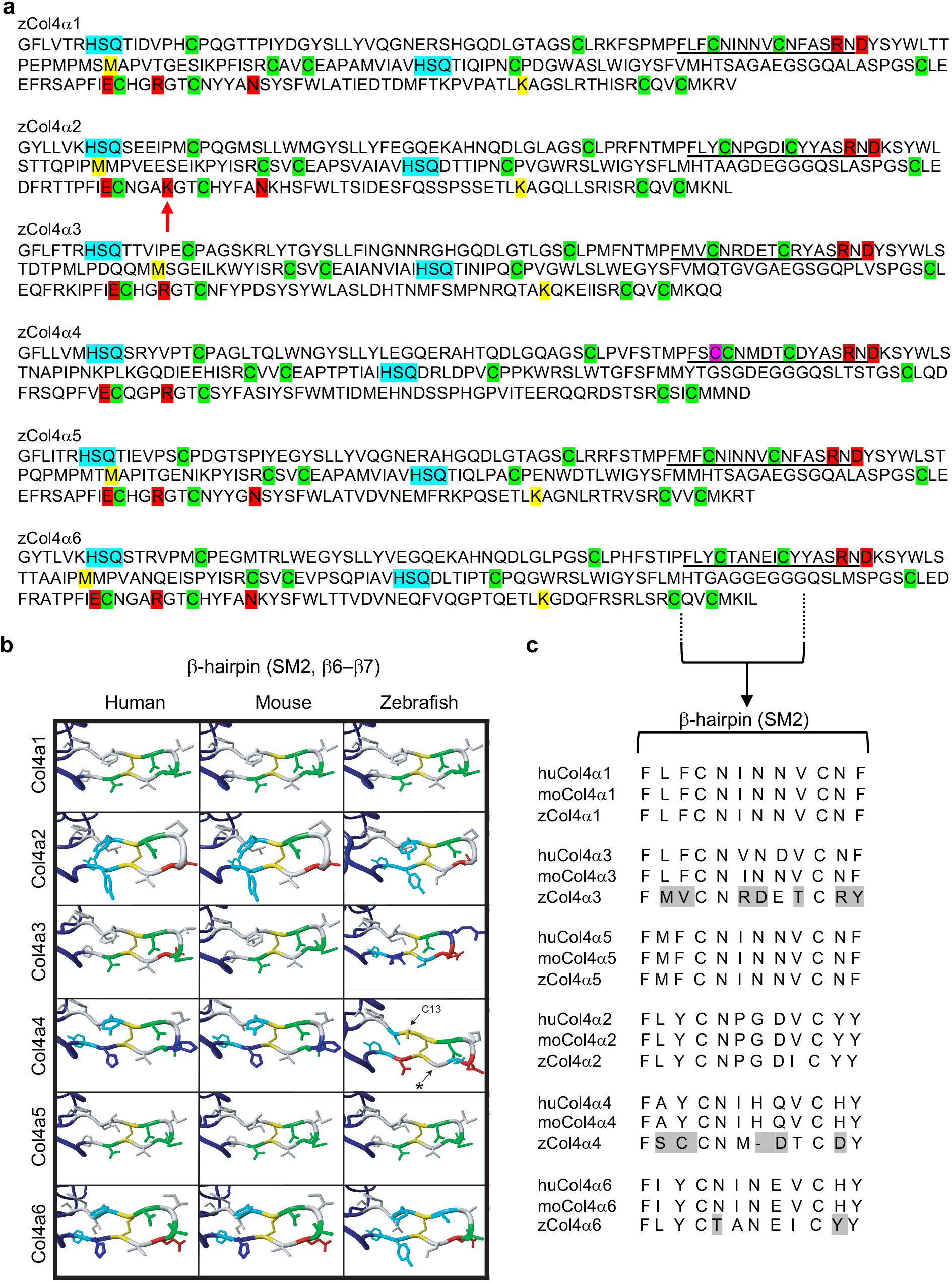
Structure of the NC1 domain of zebrafish collagen IV α-chain proteins. **a**. Sequence comparison of the zebrafish α1-α6 NC1 domains. Cyan: HSQ tripeptide that begin each NC1. Green: twelve conserved cysteine residues. Magenta: additional cysteine (Cysteine 13, Cys-13) in zCol4α4. Red: residues involved in salt bridges formed between the R76 D78, E175-R179, and N187. The red arrow points to the R179K substitution in zCol4α2. Yellow: M93 and K211 residues involved in the peroxidasin-catalyzed sulfilimine bond formation. **b**. Representation of the 3D molecular remodeling of human, mouse and zebrafish NC1 domain-hairpin SM2 subdomain. The additional cysteine (C13) in NC1-hairpin of zCol4α4 is indicated by the arrow. The asterisk points to the area where the zebrafish protein backbone is shorter than the human and mouse homologues. **c**. Amino acid sequence alignment for human, mouse, and zebrafish NC1 domain, highlighting key residues in the SM2 (hairpin 6–7) subdomain that indicates less conservation between the α3, α4 and α6 chains compared to the α1, α2 and α5 chains.

In contrast with other collagen IV α chains, the zebrafish α4 chain has two important divergent features. First, the zebrafish collagen IV α4 chain contains an additional cysteine immediately before the third cysteine residue (Cys-3) in the NC1 subdomain I, which we refer to as cysteine-13 (Cys-13; **Fig. 3a**, magenta). The Cys-3 to Cys-4 disulfide bond forms a β-hairpin structure in the SM2 subdomain (6–7) in the NC1 domain that inserts into the SM3’ subdomain of an adjacent NC1 domain [8, 54–56]. Three-dimensional modeling analysis reveals that the sulfur atoms of Cys-3 and Cys-13 are a mere 7.4 angstroms apart, and such extreme proximity makes Cys-13 a plausible alternative binding partner with Cys-4. The omission of one amino acid residue makes the −hairpin in SM2 of the zebrafish α4-chain shorter than in other species; this deletion, coupled with the natural flexibility in the peptide backbone of the domain (asterisk, **Fig. 3b**), makes Cys-13 a plausible alternative binding partner for Cys-4. *In silico* analysis with Polyphen-2 indicates that the Cys-13 substitution in human Col1a1 is likely damaging (score 1.000). We therefore hypothesize that Cys-13 in the zebrafish α4-chain alters the structure of the β-hairpin region of NC1 domain, and potentially allows for distinct interactions with other α chains and formation of novel protomer (**Fig. 3b-c**) [57]. Second, the zebrafish α4 chain lacks the M93 and K211 residues required for the peroxidasin-catalyzed sulfilimine bond formation that stabilizes end-to-end protomer dimerization [10]. Such deficit would prevent the covalent linking of α4 containing-protomers at the C terminus. These results suggest that zebrafish collagen IV networks containing the α4 chain may be weaker than their human homolog.

### Unique temporal expression patterns of type IV collagen chains in zebrafish embryologic development

To gain further insight into the biosynthesis of collagen IV protomers in zebrafish embryogenesis, we performed RT-PCR on cDNA isolated from zebrafish embryos at 13 distinct developmental time points. The results show three distinct expression patterns: temporally unrestricted expression, post-gastrulation expression, and late-onset expression (**Fig. 4a**). Expression of α1 and α2 chains mRNA was not restricted to a particular developmental stage, and the synchronous expression patterns of the α1 and α2 chains supports that zebrafish synthesize the α1α2α1 (IV) protomer throughout embryogenesis (**Fig. 4a**). In addition, transcription of genes encoding the α5 and α6 chains was synchronously detected in post-gastrulation embryos, suggesting that synthesis of the α5α6α5 (IV) protomer does not occur in the pre-gastrulation stages of embryogenesis (**Fig. 4a**). Finally, in late stage of embryos, when there was also a high level of α5 chain gene transcription, transcription of α3 and α4 chains was observed, supporting that the possible synthesis of the α3α4α5 (IV) protomer in later stages of zebrafish embryonic development (**Fig. 4a**). Our results support that the zebrafish embryo expresses the Col4 chains, suggesting the possible synthesis of the three collagen IV protomers known in mammals.

**Figure 4:**
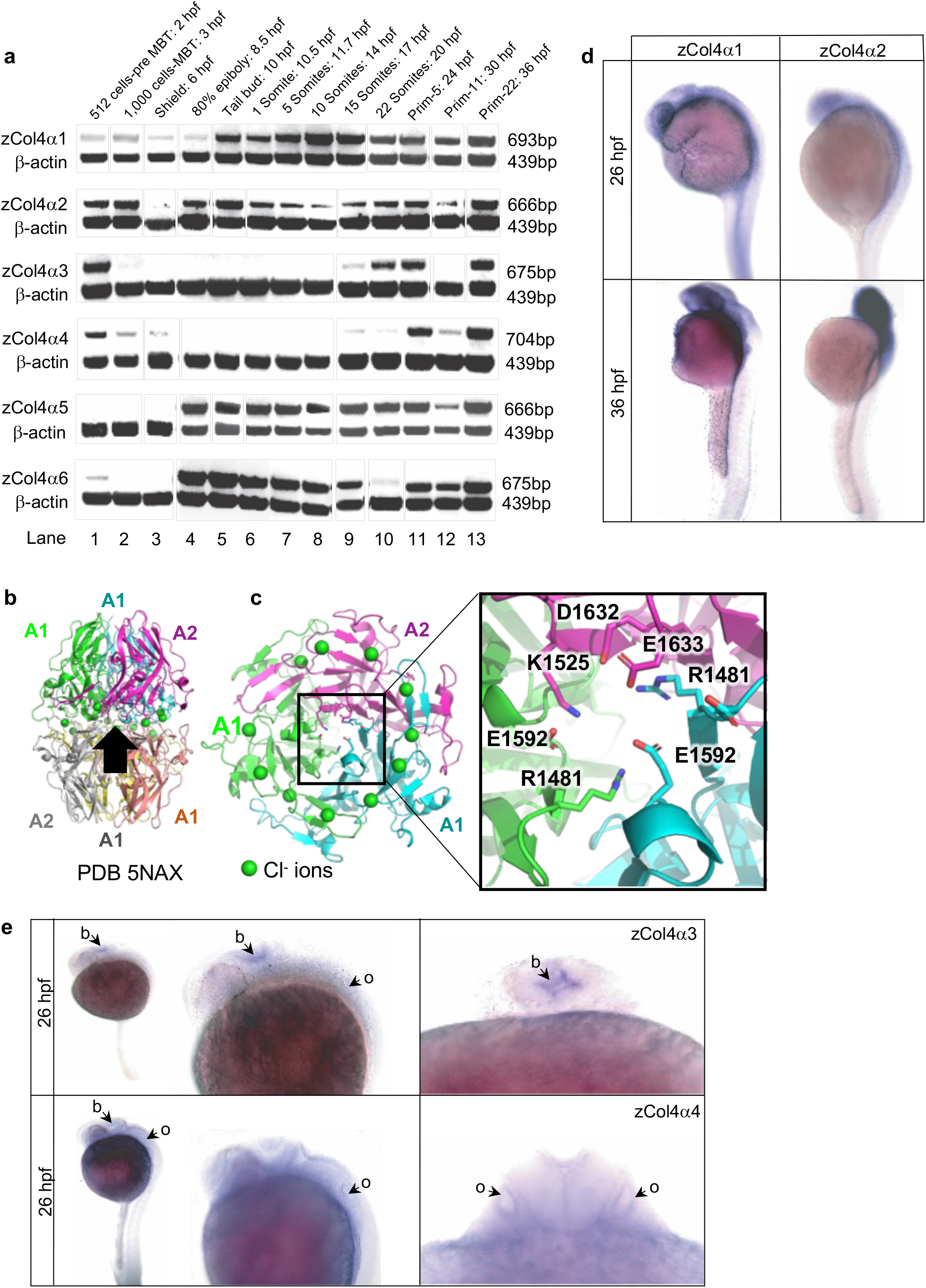
Embryonic expression of the six collagen IV α chains in zebrafish. **a**. RT-PCR products for the zebrafish type IV collagen α chains at the indicated developmental stages (top band) range in size from 666 bp to 704 bp. The β-actin control band is shown below. Three expression patterns are noted: developmentally unrestricted (α1 and α2 chains); post-gastrulation (α5 and α6 chains); and late-onset expression (α3 and α4 chains). Expression before 6 hours post fertilization (hpf) stage is maternal. **b**. Cartoon schematic of NC1 heterohexamer of α1 and α2 NC1 chains (PDB 5NAX). Arrow shows viewpoint for c. **c**. Bottom-up view of heterotrimer. Inset is close-up view of salt bridges that help form the heterotrimer interface. In the α1 and α2 chains, a glutamate (E1592 in α1 and E1633 in α2) forms a salt bridge with a basic residue (R1481 in α1 and K1525 in α2). **d**. In situ hybridization for zebrafish α1 and α2 chains at 26 hpf and 36 hpf. **e**. In situ hybridization for zebrafish α3 and α4 chains at 26 hpf, dorsal and lateral views. The α3 and α4 chain are localized in the developing brain (b), with α4 chain also detected in the otic vesicle (o). hpf, hours post fertilization, MBT, mid-blastula transition.

We also observed that the α1 and α4 chains are synchronously expressed in the absence of other chains at the shield time point (**Fig. 4a**, lane 3). In addition, there is no detectable α3 chain expression despite persistent expression of α4 at the Prim-11 stage (**Fig. 4a**, lane 12). This raises the possible existence of novel protomer involving the α4 chain other than the α3α4α5 (IV) protomer at this stage, such as α1α4α1. In zebrafish, expression of proteins before 6 hours post hybridization (hpf) stage is maternal. Only the α5 chain is not maternally contributed, which also supports that the α3- and α4-chains could assemble protomers with other composition than the α3α4α5 (IV) protomer. Though based on transcriptional evaluation only, our findings suggest distinct type IV collagen network composition may exist in the zebrafish development. Given that an α1-α4 heterohexamer is not formed in human, we postulated that mutation in zebrafish α4 allow for this formation. Targeting regions identified to be critical for oligomerization [58], we identified any changes in zebrafish α4 NC1 that made it similar to α1 and α2 NC1. Although there were several mutations that may render zebrafish α4 to be similar to α2, only one change made it similar to both α1 and α2 in one region, SM1’ (structural motif SM1’: hairpin φ33’– φ34’ [58]) (**Fig. 2**). Examination of the α1/α2 heterohexamer structure from human placenta [55] revealed that the mutation in zebrafish α4 to glutamate (E1589) would make it analogous to E1592 and E1633 in α1 and α2 respectively (**Fig. 4b-c**). These analyses raise the possibility for α4 chain containing (IV) protomer that include either the α1 or α2 chain.

### Organ-specific expression patterns of type IV collagen chains in zebrafish development

To define the anatomic position of collagen IV networks in zebrafish development, we performed *in situ* hybridization for each of the six collagen IV α-chain transcripts in the zebrafish embryo at 26 hpf and 36 hpf. Consistent with a recent study [59], our results show that the α1 and α2 chains were transcribed ubiquitously (**Fig. 4a, d**). Although the caudal expression level was low, no specific organ displayed a differential distribution of α1 or α2 chain. The α3 and α4 chains, however, displayed organ specific expression, with transcript accumulation in the brain (b) and otic vesicle (o) at 26 hpf (time point of maximal intensity; **Fig. 4e**). At 36 hpf, α6 chain expression in the otic vesicle, gills (g), and pronephric ducts (pn) could offer support for the synthesis of the α5α6α5 protomer in these tissues (**Fig 4a, Fig. 5a**).

**Figure 5:**
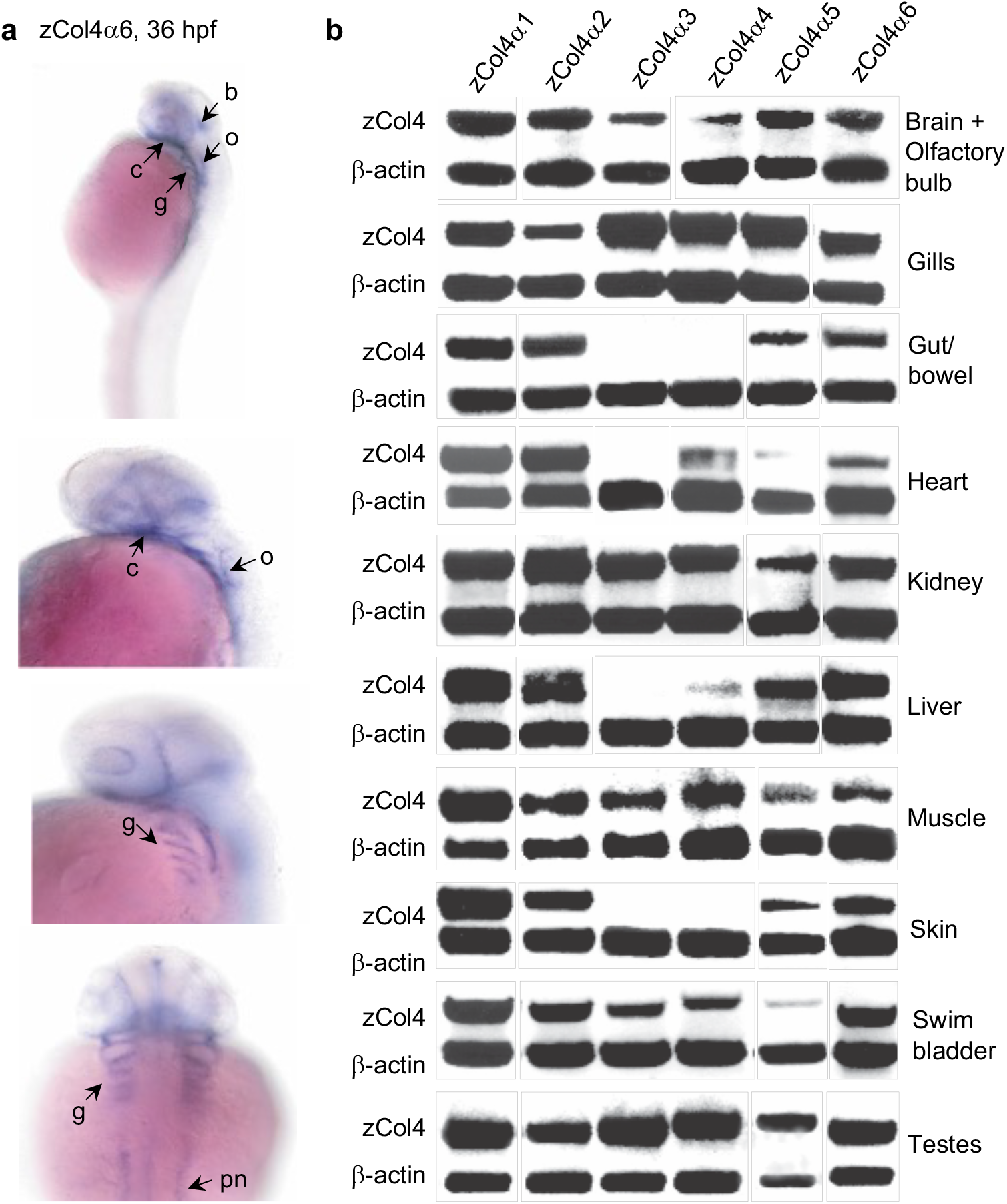
Tissue distribution of the six collagen IV α chains in zebrafish development. **a**. In situ hybridization for zebrafish α6 chains at 36 hpf in the brain (b), otic vesicle (o), gills (g), developing cardiac tissue (c), and the pronephric duct (pn) of the future kidney. **b**. RT-PCR products for the six α chains are shown above the positive control β-actin. All six α chains are detectable in brain, gills, kidney, skeletal muscle, swim bladder and testes. Gut/intestine and skin express the α1, α2, α5, and α6 chains only. Liver and heart both display expression of all α chains except the α3 chain.

In humans and mice, organs such as intestine and skin are strictly composed of the collagen IV α1α2α1 and α5α6α5 protomers, yet other organs display a developmental restricted expression of specific protomers [35, 60, 61]. For example, during human nephrogenesis, the GBM is initially comprised of the collagen IV α1α2α1 protomer. As the glomerulus matures, the GBM collagen IV undergoes an isoform switch from the α1α2α1 to the α3α4α5 protomer [62, 63]. Our results show that the expression of the more specialized α3α4α5 protomer was not detected by *in situ* hybridization in many of the zebrafish organs during embryonic development. We posit that the absence of the α3α4α5 protomer in the developing organs of the zebrafish in its embryonic development could reflect a similar post-embryonic stage isoform switching mechanism. To test this hypothesis, we performed RT-PCR for the six collagen IV alpha chain transcripts in distinct organs in the adult zebrafish. The α1, α2, α5 and α6 chains were ubiquitously expressed in the adult zebrafish (although α5 chain expression was low in the cardiac tissue), but the α3 and α4 chains exhibited a more restricted expression pattern (**Fig. 5b**). The α3 chain was expressed in the brain/olfactory bulb, gills, kidney, muscle, swim bladder, and testis, but was absent in the gut, heart, liver and skin (**Fig. 5b**). The α4 chain had a similar expression pattern, but it was also expressed in cardiac and hepatic tissue (**Fig. 5b**). These results support a protomer isoform switch in the post-embryonic zebrafish development, similarly to what is observed in mammals. The α4 chain expression in the adult cardiac tissue, concurrent with absent α3 and low α5 chain expression, indicates a more promiscuous α4 chain expression in zebrafish compared to human, and supports potential formation of novel protomers composition.

### Conserved anti-angiogenic function of the zebrafish collagen IV α3 NC1 domain, zTumstatin

Given that Tumstatin, the NC1 domain of collagen IV α3 chain, is a known anti-angiogenic cleavage product of Col4α3, we wondered whether zebrafish α3-chain NC1 domain, zTumstatin, would also exert anti-angiogenic functions, similarly to its human homolog [42, 43, 64]. To test this hypothesis, we produced recombinant zTumstatin using a eukaryotic expression system, as previously described [46]. Human umbilical vein endothelial cells (HUVEC) proliferation was inhibited in a dose-dependent manner by eukaryotic zTumstatin (**Fig. 6a**). zTumstatin also inhibited HUVEC migration in a Boyden chamber assay (**Fig. 6b**). The anti-angiogenic property of zTumstatin was also tested in vivo in a Matrigel™ plug assay. Subcutaneous Matrigel™ plugs, containing bovine FGF and VEGF, and supplemented with or without zTumstatin, were implanted in C57BL/6 mice. Control plugs (no zTumstatin) demonstrate robust angiogenesis, whereas in zTumstatin-containing plugs showed minimal levels of angiogenesis (**Fig. 6c**). Microscopic angiogenesis, evaluated by H&E staining of Matrigel™ plugs, also revealed suppressed angiogenesis (54% reduction) in zTumstatin-containing plugs compared to control (**Fig. 6d-e**). These results support antiangiogenic function of the zTumstatin.

**Figure 6:**
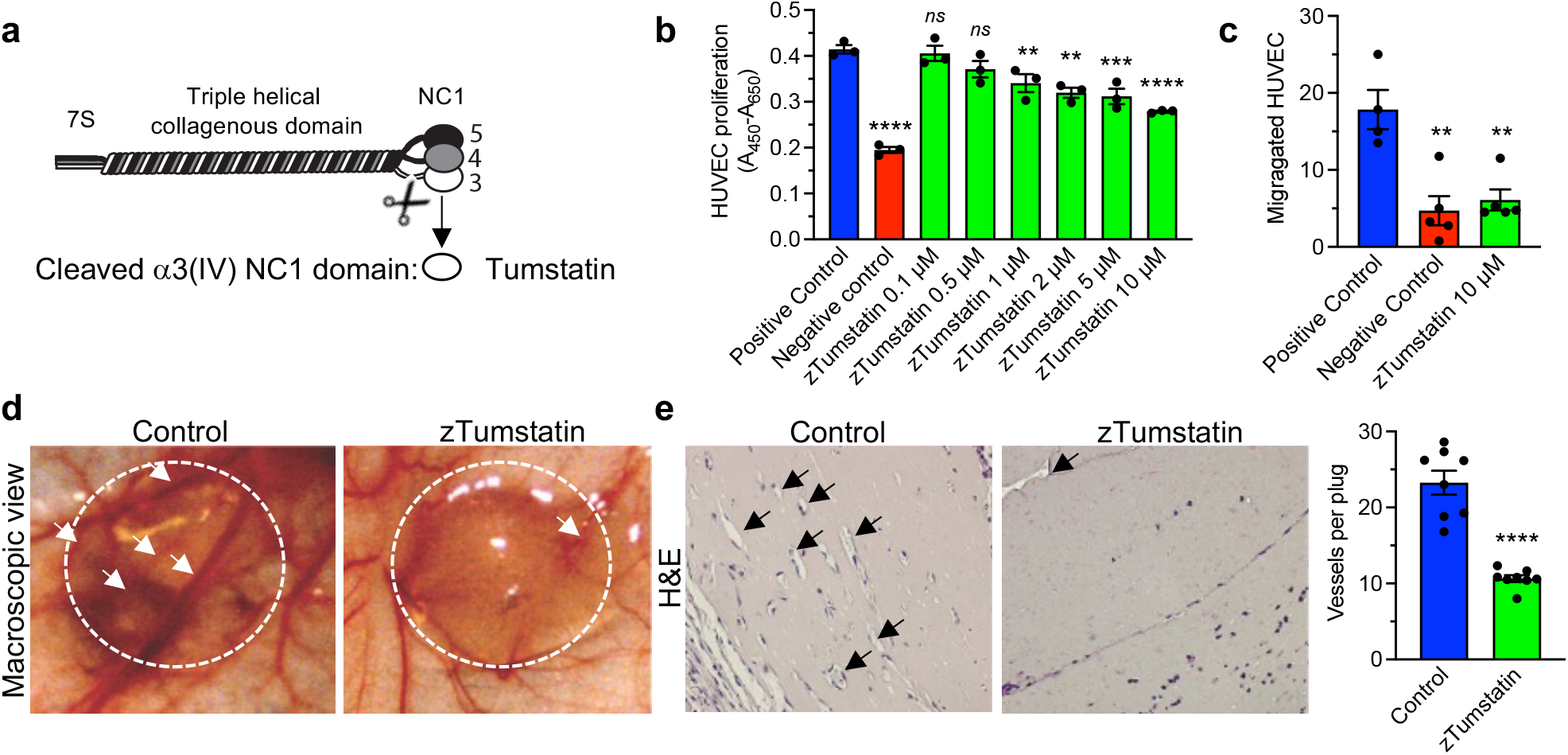
Recombinant zTumstain, the NC1 domain of the α3 chain of type IV collagen, shows conserved antiangiogenic activity on human endothelial cells. **a**. Schematic depiction of Col4 α3/α4/α5 protomer and cleavage site releasing Tumstatin (α3 NC1 domain). **b**. Relative HUVECs proliferation when treated with the indicated concentration of eukaryotically-produced zTumstatin, n = 3 wells per condition. Positive control: 10% FBS, Negative control: 0.1% FBS. **c**. HUVECs migration in a Boyden-chamber assay, n = 4-6 wells per condition. **d**. Representative macroscopic image of control and zTumstatin Matrigel™ plug (encircled) and blood vessels (arrows). **e**. Representative H&E section of control and zTumstatin Matrigel™ plug (arrows point to blood vessels) and quantification of averaged blood vessels per 200x field per plug, n = 8 plugs per group (2 plug per mouse, 4 mice per group). b-c: One-way ANOVA, e: unpaired two-tailed t test, *ns*: not significant, ** p<0.01, *** p<0.001, ****p<0.0001.

## Discussion

Collagen IV exists in all metazoans, including sponges [18]. Type IV collagen has pivotal function in shaping tissue and organs, transducing signaling pathways and maintaining normal development in multicellular organisms [3, 18]. Zebrafish, as one of the lowest vertebrates known to express the six collagen IV homologous to those in humans, is a powerful model system to study conserved function of collagen IV and therefore collagen IV-related disease. The evolution of the type IV collagen genes includes the initial tandem duplication of *Col4a1* or *Col4a2* and head-to-head organization of the *Col4a1-Col4a2* gene pair and a shared bi-directional promoter. The pair possibly duplicated onto distinct chromosomes, resulting in a similar head-to-head organization of the *Col4a3-Col4a4* and *Col4a5-Col4a6* pairs; and this gene arrangement is conserved between zebrafish, mouse, and human. This may explain the increased similarities in the NC1 domains, conserved to preserve specific network assembly, of the α3 chain with α1 and α5 chain, and the increased similarities in the α4 chain with the α2 and α6 chains [18, 65]. The comparison of the sequence of -hairpin SM2 subdomain and SM3’ subdomain therein also supports this interpretation.

We also show the conserved twelve cysteine residues required for maintenance of NC1 domain structure in all six zebrafish α chains when compared to human NC1 domains. The residues involved in the formation of the chloride-salt-bridge-mediated sulfilimine bond, through a conformational change of NC1 domain, are conserved in all six collagen IV α chains in zebrafish. In addition, the enzymes that catalyze collagen IV protomer cross-linking in the NC1 and 7-S domains, peroxidasin and lysyl oxidase-like 2 [11, 56, 66], are also conserved. We note that the zebrafish α4-chain NC1 domain lacks M93 and K211 residues and contains an additional cysteine residue. The M93 and K211 residues are joined by a sulfilimine bond required for linking adjacent protomers [10, 55], suggesting distinct adjacent protomer interaction for the zebrafish α4-containing protomer. 3D remodeling in our study also supports that the additional cysteine residue alters the zebrafish α4-containing protomer interaction with other protomers.

Our study also offers a comprehensive analysis of the temporospatial expression pattern of each α chain throughout zebrafish development at the RNA level. Our results support that zebrafish generates the collagen IV protomers known to be synthesized in humans: α1α2α1, α3α4α5 and α5α6α5. Similarly, in mice, spatial and temporal distribution of collagen IV isoforms in the developing eye also reveals these protomers [67]. In our study, we note that the transcriptional pattern of α1 and α2 chain is widely distributed, whereas which of the α3-α6 chain exhibits a restricted, organ specificity pattern, consistent with recent findings [59]. Synchronous expression of α3-, α4- and α5-chains in the brain suggests that α3α4α5 (IV) protomer is synthesized there at an early time point, which agrees with the results of a recent study highlighting the importance of this protomer in zebrafish neural development [65]. Furthermore, synchronous expression of these α chains in the otic vesicle and pronephros support an evolutionary conserved anatomic specificity for α3α4α5 synthesis, given that this protomer is the major component of the cochlear and glomerular basement membranes in humans. The chains expression enabling α3α4α5 network synthesis is not detectable in these organs until late time points, which supports a similar, evolutionary conserved, collagen IV isoform switching event in these tissues [62, 63].

Our findings unexpectedly yielded complementary evidence (homology and temporospatial expression) that the zebrafish collagen IV protomer repertoire may include novel protomers. For instance, zebrafish α4 chain was synchronously expressed with the α1 chain in the absence of α2, α3 and α5 chains at the shield time point and in cardiac tissue of adult zebrafish, wherein α3 and α5 chain expression were lacking or were minimal. These findings suggest the potential existence of a putative α1α4α1 protomer. The zebrafish α4 chain NC1 domain containing an additional cysteine residue, which is predicted to lead to structural changes of the -hairpin loop, may enable additional diversity of protomers [54–56]. Lack of M93 and K211 residues and an additional cysteine residue may trigger distinct adjacent protomer interaction for the zebrafish α4-containing protomer. Collectively, our results suggest that apart from α3α4α5, other potential collagen IV protomers (α1α4α1) may present in zebrafish. Although, our analyses rely on gene transcription findings, we previously showed protein-level distribution of collagen IV α3 [46]. Collagen IV α3 protein expression in the adult zebrafish is in the gills and kidney [46], which is in agreement with our RNA analyses. Collagen IV α4, using protein trap insertional mutagenesis was detected in the caudal vascular plexus and myotomes of the developing zebrafish [68]. Future studies are needed to evaluate zebrafish α chain interactions at the protein level.

Finally, we report on the conserved function of the zebrafish collagen IV α3 chain NC1 domain (zTumstatin). Similar to its human ortholog (Tumstatin), zTumstatin presents with anti-angiogenic activity in vitro and in vivo. The epitope that is targeted by autoantibodies in Goodpasture syndrome is however not conserved [46]. This suggests a distinct immune system maturation and regulation of the zebrafish compared to human, with respect to the autoimmune response to collagen IV α3 chain in Goodpasture syndrome [69]. Our studies reveal evolutionary conservation of collagen IV NC1 domain between zebrafish and humans at the molecular and functional level and, together with other reports, made advances in understanding collagens by using model organisms [70].

## Experimental Procedures

### NC1 Domain Molecular Analysis and 3D rendering

Amino acid sequences were obtained from UniProt database. The color scheme used for alignments is from the algorithm in Clustal Omega and MView. Each amino acid is colored when it meets the criteria specific for the residue type. This analysis is conducted by Clustal X and viewed by Jalview. The 3D modeling of the structures of the [(α1)2(α2)]2 NC1 hexamers from human placenta basement membranes (Protein Data Bank ID 1LI1) were rendered using the molecular graphics visualization program YASARA (YASARA Bio-sciences) and analyzed to generate modeled structures using the molecular graphics software InsightII (Accelrys). Percentage of homology was calculated by multiplying the query coverage and percent identity of each comparison with usage of NCBI blast tool.

### Animal Care

AB and Tu strain zebrafish were bred and maintained as described [46]. C57BL/6 mice (Charles River Laboratories) were used for the *in vivo* angiogenesis assay. Animal studies were reviewed and approved by the institutional animal care and use committee at Beth Israel Deaconess Medical Center and Boston Children’s Hospital.

### RT-PCR Determination of Embryonic and Adult Expression Pattern

At each of the 13 developmental stages described therein, total RNA was isolated, and cDNA was generated from pools of 25 embryos. For adult AB strain, ten distinct organs were isolated, placed in TRIzol™ (Invitrogen) reagent and immediately homogenized. Generation of cDNA was performed using Superscript II™ reverse transcriptase (Invitrogen) and oligo dT primers (Invitrogen). RT-PCR primers (below) were designed to regions more unique than those used to clone the NC1 domains and using the NCBI BLAST program to assure that none of these primers could cross-amplify any other zebrafish type IV collagen α chains.

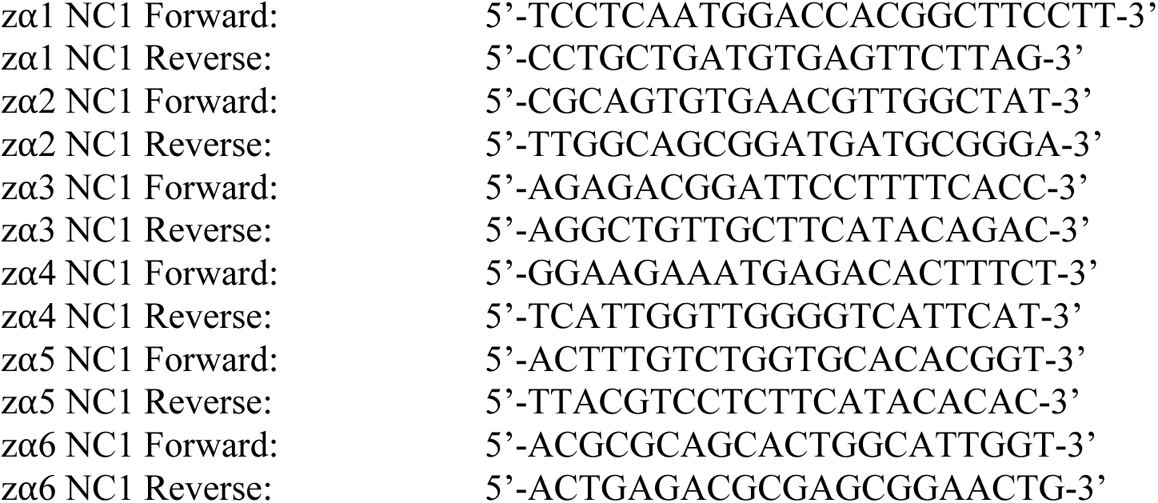

### Probe Generation and Whole Mount *in-situ* Hybridization of Embryos

Whole-mount *in situ* hybridization was performed as described [71]. Digoxygenin-labeled antisense RNA probes were synthesized using a DIG RNA Labeling NTP mix (Roche) and either SP6 or T7 RNA polymerase (Invitrogen). Primers used for generating probes to amplify the 3’-untranslated regions of type IV collagen transcripts (α1-α6) in zebrafish are:

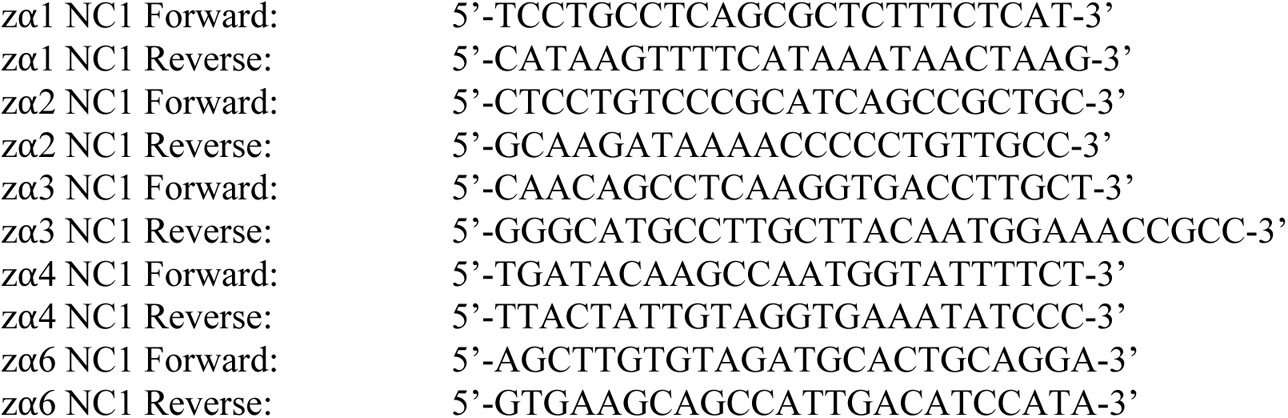

### Recombinant Protein Production

A cDNA encoding zTumstatin (zebrafish Tumstatin), was subcloned into the pcBFT vector containing a FLAG tag. The vectors were then transfected into human embryonic kidney cells (HEK293) with Fectin-system (Invitrogen). Protein from culture supernatant was isolated as previously described [46, 72].

### In Vitro Cell Proliferation Assay

Human umbilical vein endothelial cells (HUVEC) were obtained from Lonza and cultured in EGM-2 cell culture medium (Lonza). Assays were carried out in 96-well plates, with 6,000 cells per well and using WST-1 reagent (Roche). Absorbances were measured at 450nm and 650nm on a Versamax turntable microplate reader (Molecular Devices). Three independent wells were measured for each condition and the data presented as absorbance (A) values at 650nm subtracted from 450nm (A_450_ - A_650_). Positive control: 10% FBS; Negative control: 0.1% FBS. HEK293 produced zebrafish Tumstatin was supplemented to 10% FBS at the indicated concentration.

### Migration Assay

HUVEC (5,000 cells/ well) were seeded onto the top chambers of a 48-well Boyden chamber (Neuro Probe Inc) in the presence of un-supplemented media or media supplemented with 10 μM zTumstatin. HUVEC were separated from bottom wells containing EBM culture media (Lonza) supplemented with 0.1% FBS and 40 ng/ml of recombinant VEGF (R&D). The negative control did not contain VEGF in the bottom well. Top and bottom wells were separated by 8 μm polycarbonate membranes (Neuro Probe Inc). Membranes were fixed and stained using the PROTOCOL HEMA 3 Stain Set (Fisher Scientific) after removing all cells from the top side of the membrane. The number of migrated cells was scored under a 20x objective by light microscopy. Each well was evaluated with 2 to 4 images captured per well. For each group, 4 to 5 wells were evaluated.

### Matrigel™ Plug Assay

Matrigel™ (BD Biosciences) was mixed with 20 units/ml heparin (Pierce), 50 ng/ml bFGF and VEGF (R&D), and 10 μmol/L zTumstatin. Control plugs did not contain zTumstatin. The Matrigel™ mixture was injected subcutaneously in C57Bl/6 mice, 2 plugs per mouse. After 7 days, the mice were sacrificed, and the Matrigel™ plugs were removed and fixed in 10% buffered formalin. The plugs were then embedded in paraffin, sectioned, and stained by hematoxylin and eosin. Sections were examined by light microscopy, and the number of blood vessels counted from five or more images taken for each plug, and the number of vessels counted was averaged for each plug. A total of 8 plugs from 4 mice were analyzed in each group, statistical analyses were performed on plugs.

### Quantification and statistical analysis

GraphPad Prism software was used for graphic representation and statistical analysis. One-way ANOVA method was used in **Fig. 6b-c**, compared to positive control. Two-tailed unpaired Student’s t test was used in **Fig. 6e**. The data is presented as the mean +/- the standard error of the mean. Significance of statistical tests is reported in all graphs as follows: ****, p<0.0001; ***, p<0.001; **, p<0.01.

## Abbreviations

NC1 domain: non-collagenous 1 domain
GBM: glomerular basement membrane
hpf: hour post fertilization

## Acknowledgements

This study was partially funded by a research grant from Emerald Foundation, NIH research grant DK55001 to Raghu Kalluri, Cancer Prevention and Research Institute of Texas, and research funds from the MDACC and Beth Israel Deaconess Medical Center for the Division of Matrix Biology. The 2008 summer sabbatical of MS was funded by SSMF and the Swedish Society for Medicine.The authors would like to express our gratitude to Noelle Paffett-Lugassy and Caroline Burns for guidance in isolating the numerous adult zebrafish tissues.

## Author Contributions

VSL, ST, JD and MS reviewed the data, figures, and revised the manuscript. BAM, HS, LX, LIZ, and JT performed or supervised experiments. JLA performed the initial homology analysis. VSL, ST, JD, BAM prepared the figures. RK oversaw the scientific design of the project and edited the manuscript.

